# Ozboneviz: An Australian Precedent in FAIR 3D Imagery and Extended Biodiversity Collections

**DOI:** 10.1101/2024.12.04.626889

**Authors:** Vera Weisbecker, Diana Fusco, Sandy Ingleby, Ariana B J Lambrides, Tiina Manne, Keith Maguire, Sue O’Connor, Thomas J Peachey, Sofia C Samper Carro, David Stemmer, Jorgo Ristevski, Jacob D van Zoelen, Pietro Viacava, Adam M Yates, Erin Mein

## Abstract

Billions of specimens are in biodiversity collections worldwide, and this infrastructure is crucial for research on Earth’s natural history. Three-dimensional (3D) imagery of specimens is an increasingly important part of the ‘Digital Extended Specimen’ network of metadata. Open access, high fidelity 3D imagery of biodiversity specimens improves researcher efficiency, equity and increases public engagement with collections. We introduce Ozboneviz, an open access collection of FAIR (Findable, Accessible, Interoperable, Reusable) 3D imagery aiming to enhance research capacity in Australasian vertebrate skeletal morphology. Ozboneviz is an Australian test case demonstrating the feasibility of creating multi-institutional, FAIR 3D biodiversity imagery collections. We outline project design, challenges, and use by the international research community. We then discuss the urgent need for investment in infrastructure and curatorial support to progress the digitisation of Australian biodiversity collections in a way that maximises stakeholder benefit and facilitates 3D data discoverability and retrieval.

## Introduction

Biodiversity collections have been the mainstay of research in areas such as evolutionary biology, palaeontology, archaeology, taxonomy and ecology (Beck et al. 2022, Hilton et al. 2021, Holmes et al. 2016, Lyman 2019, McLean et al. 2016, Meineke et al. 2018). They also have critical industrial applications such as for pest and invasive species management, biosecurity, agriculture, and conservation, as well as playing a major role in public and tertiary education (Atlas of Living Australia et al. 2023, Ball-Damerow et al. 2019, Cook et al. 2014, Lyal et al. 2008, National Academies of Sciences and Medicine 2020, Suarez and Tsutsui 2004). Accessing the three-dimensional (3D) morphology of biodiversity specimens is thus a core requirement across the academic, commercial, and government sectors. Although curation of physical vouchered specimens remains crucial, high quality 3D imagery can facilitate education and research programs simultaneously and efficiently.

For many stakeholders, access to biodiversity collections is inefficient, costly when travel to collections is required, and potentially damaging to specimens due to repeated handling (Davies et al. 2017, Hipsley and Sherratt 2019, Johnson et al. 2023, Page et al. 2015). Public engagement with these mostly taxpayer-funded research collections is also limited to curated exhibition initiatives, principally in museums (Blackburn et al. 2024, Boyer et al. 2016, Nelson and Ellis 2019, Powers et al. 2014). Increased access to these collections improves research efficiency, reduces inequities in the research community, and deepens and diversifies public engagement in this important infrastructure (Atlas of Living Australia et al. 2023, Blackburn et al. 2024, Cook et al. 2014, Drew et al. 2017, Hedrick et al. 2020, Hipsley and Sherratt 2019, Johnson et al. 2023, Lendemer et al. 2020, Nelson and Ellis 2019).

More equitable specimen access can be, in part, achieved through high-fidelity 3D imagery that acts as a digital ‘avatar’ of the physical specimen (Hipsley and Sherratt 2019). An ongoing revolution in 3D digital imaging has made the creation of these avatars increasingly affordable (Boyer et al. 2016, Davies et al. 2017, Nelson and Ellis 2019, Rowe and Frank 2011). Modalities such as magnetic resonance imaging (MRI), X-ray computed tomography (CT) are often more informative than the physical examination of the vouchered specimen because they can reveal internal or small-scale features that are difficult to observe with the naked eye or without destructive sampling (Blackburn et al. 2024, Hilton et al. 2021, Kimura 2023). 3D imagery can be used to retain information prior to destructive sampling or insure against total data loss during disasters (Crown v Van Leeuwen 2007, Escobar 2018, Tyler et al. 2023). It can be easily and quickly disseminated (Blackburn et al. 2024, Boyer et al. 2016, Davies et al. 2017, Hipsley and Sherratt 2019) and, when curated well, 3D imagery is an important part of the Digital Extended Specimen, (Hardisty et al. 2022, Lendemer et al. 2020, Webster 2017) the connected web of data and metadata associated with a single specimen. Use of 3D imagery is rapidly increasing in biodiversity research (Čerňanský and Syromyatnikova 2019, Dong et al. 2022, Early et al. 2020, Maden et al. 2023), outreach and education (Flemming et al. 2020, Gray et al. 2024, Ulguim 2018, Ward et al. 2023) and has become integral to downstream analyses of biodiversity patterns, such as (geometric) morphometrics (Evers et al. 2022, Gray et al. 2019, Navalón et al. 2022, Weisbecker et al. 2021) or finite element analysis (Cox et al. 2015, Mitchell et al. 2021, Oldfield et al. 2012).

Despite the promise of digital 3D collections, there continue to be hurdles to the roll-out of open-access 3D imagery. For example, external, primarily university-based, researchers are major generators of 3D imagery of biodiversity specimens in Australia (Weisbecker et al. 2024). Yet perceived disincentives for data sharing continue to persist, owing to the reliance on competitive research grants to fund the often labour-intensive process of digitisation and fear of being ‘scooped’ (Hipsley and Sherratt 2019). Such monopolisation of data access, in Australia and elsewhere, creates inequities in the research community that disproportionately affect early career and unaffiliated researchers and scientists from low-and middle-income countries (Boyer et al. 2016, Davies et al. 2017, Drew et al. 2017, Hipsley and Sherratt 2019). This issue is exacerbated by the limited capacity of collections to curate these 3D data, resulting in a near-total lack of institutional oversight even when researchers are willing to share their data (Atlas of Living Australia et al. 2023, Rowe and Frank 2011, Weisbecker et al. 2024). Fortunately, scientific consensus on data sharing is shifting and there is an increasing expectation from funding bodies and journals that scientific data – including 3D imagery - be made open access and FAIR (Australian Research Council 2022, European Research Council 2024, Foundation 2023, National Health and Medical Research Council 2019, Nature Portfolio Journals 2024, OECD 2007, Science Journals 2024, The Royal Society 2024, Wilkinson et al. 2016). With robust policy frameworks and user-friendly implementation, researchers will likely adopt FAIR 3D data sharing in a similar way as previously implemented with genetic data through initiatives like GenBank (Benson et al. 2015) or the International Nucleotide Sequence Database Collaboration (International Nucleotide Sequence Database Collaboration 2024).

Although FAIR research is a key goal in an equitable research culture, project-based 3D digitisation by individuals cannot progress biodiversity digitisation in a way that maximises the scientific and public benefit of collections. Digitisation initiatives are needed which prioritise the collection of 3D imagery to service the needs of diverse stakeholders – these will be termed “service collections” herein. Service collections are already well advanced in some parts of the world (Berlin 2024, Harvard University 2018, Le Bras et al. 2017, Scott et al. 2019, Smithsonian Institution 2024, Thiers et al. 2016); one of the largest and most successful single 3D imaging initiatives is the National Science Foundation (NSF) funded oVert project which provides global access to over 29,000 high fidelity 3D models representing more than 13,000 specimens from 50 institutions (Blackburn et al. 2024). In Australia, however, there is no local precedent for creating, storing or curating such a multi-institutional service collection of 3D imagery. One of the reasons for this shortfall is the lack of consensus and frameworks for addressing a variety of concerns, ranging from practical considerations such as metadata curation to intellectual property and copyright arrangements that minimise legal risk and balance the needs of institutions and collections users (Davies et al. 2017, Hipsley and Sherratt 2019, Matsui and Kimura 2022, Weisbecker et al. 2024). Furthermore, obtaining funding to support initiatives like service collections is challenging in the current Australian research funding system.

Addressing these policy and infrastructure challenges is an urgent issue because Australian biodiversity is of substantial interest internationally. Australian flora and fauna are highly endemic and an important part of the global story of evolution and diversity of life (Dickman 2018). Australia also leads the world in vertebrate extinctions (Woinarski et al. 2019) and therefore Australian collections play an especially important role as repositories of historical biodiversity data (Haouchar et al. 2016, McDowell et al. 2015, Travouillon et al. 2019, Woinarski et al. 2019). FAIR 3D imagery would provide more equitable and diverse community access (Drew et al. 2017, Johnson et al. 2023, Salomon et al. 2023) to these Australian collections while preserving the research mandates of curating institutions. It is therefore critical that planning and policies for data acquisition, storage and integration with the Digital Extended Specimen data in Australia keep pace with global developments in 3D imaging.

Here, we introduce Ozboneviz, an open access collection of high-fidelity 3D imagery of Australasian vertebrates hosted on the MorphoSource.org (Boyer et al. 2016) repository. Funded by the Australian Research Council (ARC) Centre of Excellence for Australian Biodiversity and Heritage (CABAH) the aims of our project were two-fold: 1) to create a 3D service collection that elevates research capacity into Australasian vertebrate diversity, with particular focus on larger-bodied mammals of zooarchaeological relevance. 2) to implement a multi-institutional Australian precedent addressing the challenges of providing open and FAIR 3D biodiversity imagery. In this paper we describe the project design, status, and community use of the collection. We discuss the implementation of FAIR principles to these data, the usefulness of service collections such as ours to the international research community, as well as benefits to diverse stakeholders and curating institutions. For detailed acquisition methods, please refer to the Methods supplement.

### Collection contents

At the time of writing, the Ozboneviz collection (www.morphosource.org/projects/000394988) contains 1591 3D meshes of individual skeletal elements and 46 micro-computed tomography (µCT) images, focusing on the skull and eight major elements from the appendicular skeleton for each species (Figure 1; see supplementary methods). A total of 276 individual specimens were imaged representing 189 vertebrate species. We captured 3D data using a range of modalities with a focus on structured light surface scanning (*n* = 1109) followed by µCT segmentation (*n* = 476) with a small number of meshes created via photogrammetry (*n* = 6). A total of 170 mammal species from Australia and New Guinea were digitised including 5 monotremes, 122 marsupials and 43 placental mammals (including non-native and marine mammals) (Figure 2). We also digitised eight birds, eight reptiles and three amphibians. We acquired 3D imagery of 10 extinct species (Figure 2) and digitised elements from a paratype of the northern pig-footed bandicoot (*Chaeropus yirratji* SAMA-M3971) and the holotype of golden mantled tree-kangaroo (*Dendrolagus goodfellowi pulcherrimus* AM-M.21717). In five cases, we digitised captive specimens that we deemed as important additions to the collection, because wild representatives of the species were either not available or had insufficient metadata. Organisations that contributed over 10 specimens to the project included the South Australian Museum (*n* = 121), the Australian Museum (*n* = 68), the Museum and Art Gallery of the Northern Territory (*n* = 38) and the University of Queensland (*n* = 19) (Methods Supplement and Figure 2).

**Figure 1.**
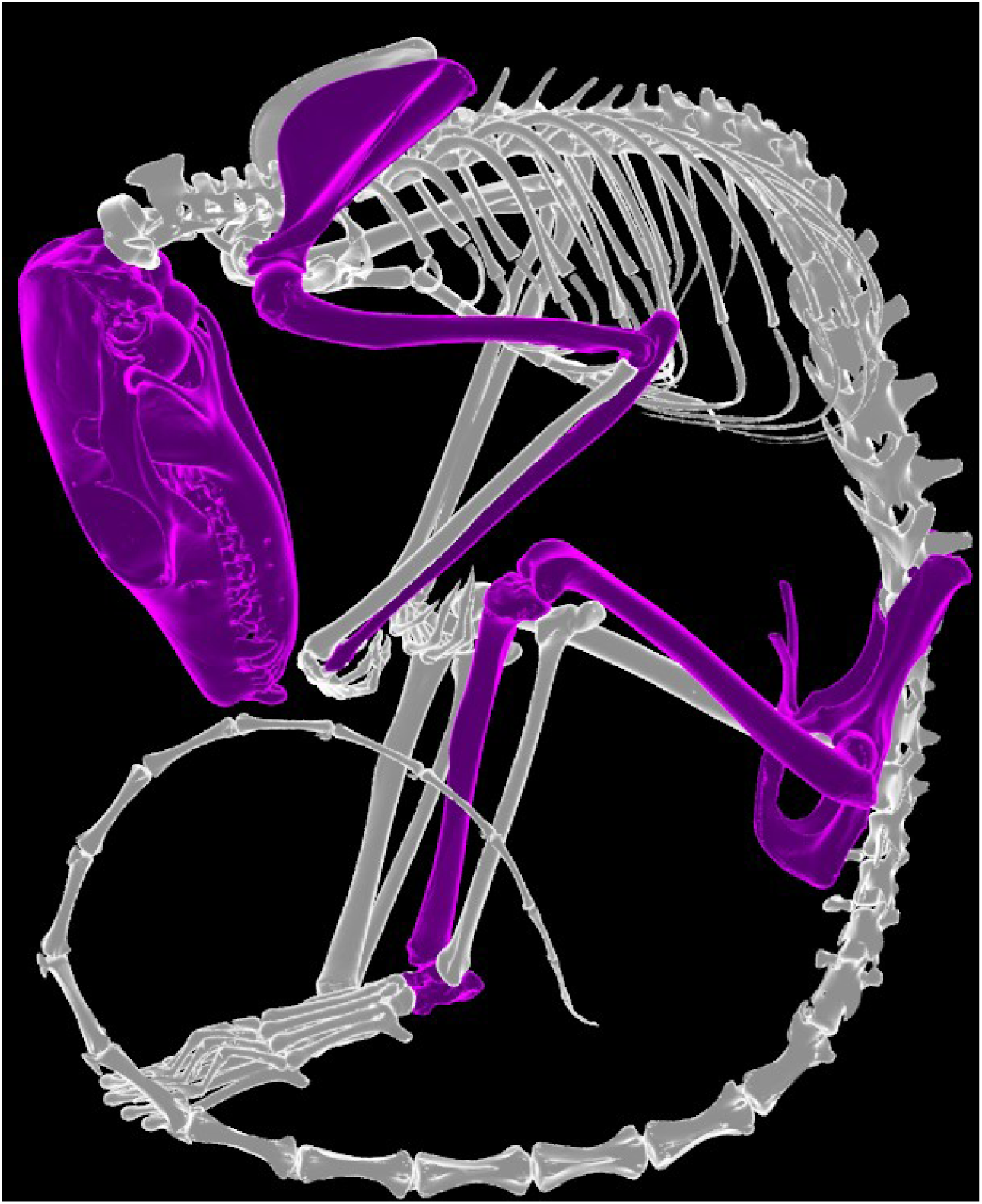
Skeletal elements from the mammal and reptile skeleton digitised by the Ozboneviz project highlighted in purple on a brush-tailed phascogale (*Phascogale tapoatafa)* specimen (SAMA-M3824).

**Figure 2.**
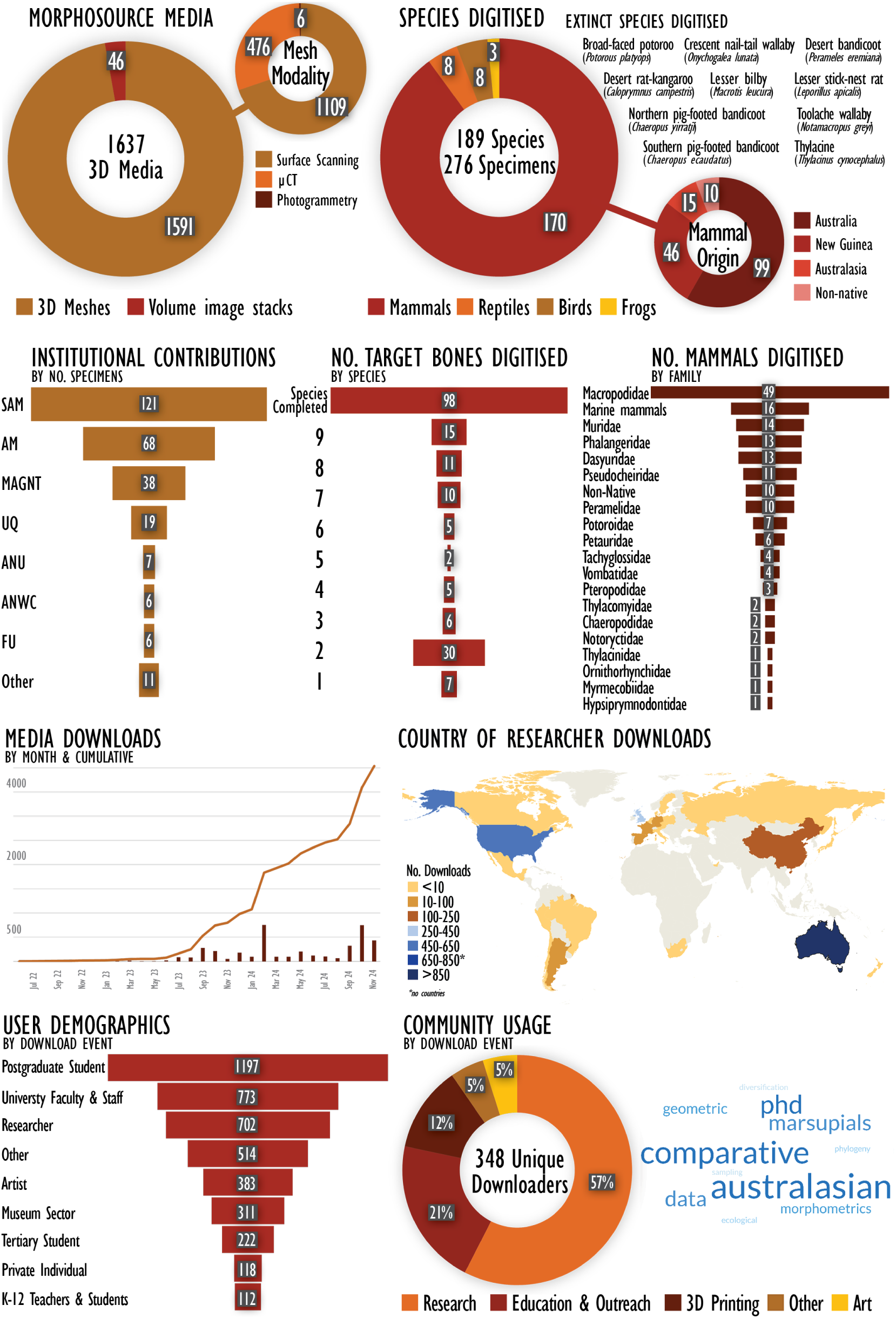
Summary of the Ozboneviz collection 3D imagery acquisition and community usage as of 26 November 2024.

### Open access, FAIR 3D imagery

We chose the NSF-funded MorphoSource platform to deposit our collection, as this was most aligned with our aim to make Ozboneviz data as FAIR (Wilkinson et al. 2016) as possible. The architecture of MorphoSource allows for connections between 3D data and specimen metadata to be maintained, as well as supporting metadata describing imagery acquisition, processing, intellectual property and copyright. It also allows data managers to authorise user access to 3D data and track data downloads, intended usage and user demographics. The platform is also becoming the preferred solution for 3D data storage by US institutions (Blackburn et al. 2024) and provides the facility for collections managers to assume control of the long-term management of 3D specimen imagery (Boyer et al. 2016), making it internationally compatible. The institutional uptake and facilitation by MorphoSource also ensure these data are likely to remain managed and accurate into the future.

FAIR data are defined as findable by human and machine, have a unique and persistent identifier, are richly described by metadata and are openly accessible with appropriate authentication protocols where necessary (Carroll et al. 2020, Hardisty et al. 2022, Sterner and Elliott 2023, Wilkinson et al. 2016). FAIR data (and metadata) should also be reusable and have clearly defined parameters for re-use. Following these requirements, researchers external to our project group have minted 83 DOIs for our 3D imagery on MorphoSource. In addition, 211 specimens are linked to an occurrence record on iDigBio, which allows the specimen metadata on MorphoSource to remain synchronised with the institutional metadata over time.

All the Ozboneviz collection are freely available to be downloaded and reused by registering a user account with MorphoSource. Conditions for reuse of these 3D data are clearly stated in the CC-BY-NC copyright policy and standard MorphoSource agreement. Users must agree to these conditions and receive a copy of the licence and usage agreement when downloading each media. Note also that copyright conditions can be changed to suit the managing collections.

### Community usage

MorphoSource captures information on the proposed use and demographics of users as a condition of downloading data. These data are self-reported by users who can select multiple options to describe themselves and their proposed data use. As of November 2024, 1232 individual models have been downloaded a total of 4044 times. The most common reported usage of these data is for research (57%) followed by education and outreach (21%) and 3D printing (12%) (Figure 2). Users who identified as post-graduate students are the largest downloaders of our data. Educators, students and university-based researchers from institutions in North America, South America, Europe, Asia, Africa and Australasia have downloaded our 3D imagery a total of 2619 times (Figure 2). A small but unexpected use of these 3D imagery has been by artists either as reference models for traditional analogue art practices (drawing, modelling etc) or for use in digital art practices. To date, the four most downloaded 3D media in the Ozboneviz collection are a cranium of a saltwater crocodile (*Crocodylus porosus*), cattle (*Bos taurus*) cranium, thylacine (*Thylacinus cynocephalus*) cranium and a New Guinea naked-backed fruit bat (*Dobsonia magna*) µCT image stack.

### Technical Challenges

We encountered several challenges in assembling the Ozboneviz collection related to specimen suitability for imaging and specimen metadata. Our project goal was to digitise eight to 10 bones per species (depending on taxonomic class) and our core acquisition modality was surface scanning of dry, skeletonised specimens. However, digitising all our target bones for each species was more challenging than anticipated, even where we were able to digitise bones from multiple individuals of the same species. Post-cranial skeletal elements are less frequently preserved in biodiversity collections than skulls or were often unsuitable for surface scanning owing to adhering soft tissue, translucency caused by bone grease or articulation with neighbouring bones. We successfully digitised all target bones for 104 species, although 50 species have only five or less bones digitised (Figure 2). We also found the completeness of specimen metadata varied widely owing to historical realities of collections management. For example, many specimens were provided as roadkill were too decayed to provide a sex, which can be important information for downstream analyses. In these cases, we prioritised specimen intactness and our ability to produce high quality 3D imagery over metadata completeness.

The extensive use of µCT for full skeleton acquisitions was not within the scope of our funding, but we were able to choose a selection of particularly rare or important species for full-body µCT acquisitions (see methods supplement for additional challenges specific to this modality). Non-native and domestic animals were also an important component of our service collection but are often not curated in Australian museum biodiversity collections. We primarily relied on university-based collections for placental mammals such as foxes, cats, dogs, horses and domestic artiodactyls. However, we found that specimen metadata in university collections was generally poorer than museum-based collections. Furthermore, most university collections are not published on biodiversity aggregators such as GBIF or iDigBio and, although most have protocols for allowing researchers access, these are not publicly discoverable.

### Permission challenges

Because of the lack of an Australian precedent, it was particularly challenging to identify appropriate terms under which specimens could be acquired and published to the satisfaction of participating institutions. The core team engaged in extensive consultation with collection representatives to discuss diverse institutional perspectives on issues such as scan ownership, intellectual property, and copyright. For this purpose, our team developed an FAQ document with support from the MorphoSource Team. Although all collections we approached were supportive of the concept of Ozboneviz, the relative novelty of the “service collection” model we adopted made it difficult to assess the risk and some collections felt unable to issue a permission for data acquisition. The South Australian Museum issued a formal permission text which became a useful basis for permissions from other collections.

### Ozboneviz as a service collection for Australian vertebrate skeletal 3D imagery

Ozboneviz has succeeded in providing a comprehensive skeletal database of Australian land vertebrates. We see it as a timely contribution in the continuously expanding landscape of 3D biodiversity data, based on the enthusiastic adoption by the scientific and wider community.

Our usage data show that the initial scope of Ozboneviz as a resource for zooarchaeological data was quickly expanded into other areas of science (such as palaeontology and evolutionary biology), public engagement, and the arts. This is an excellent example of how the availability of open-access data can have unforeseen benefits well beyond the original scope of the data collection if it is made accessible in appropriate ways (Blackburn et al. 2024, Davies et al. 2017, Lendemer et al. 2020, Suarez and Tsutsui 2004). In addition, despite its novelty, several international research efforts are using Ozboneviz data, demonstrating the almost immediate positive impact of the collection in representing Australian vertebrate diversity to the international research community.

As outlined in the introduction, one of the greatest potential benefits of open-access 3D data is the improvement in equitability of access to researchers and other stakeholders (Cook et al. 2014, Drew et al. 2017, Hipsley and Sherratt 2019). The urgency of this is particularly stark in our user statistics, with by far the greatest download activity coming from the most junior researcher cohort - postgraduate students. The appetite for these data is clearly substantial among these early-career researchers, who are often the most disadvantaged in terms of the funding, time and reputation required to individually access these specimens. In addition, nearly a quarter of our downloads are reported for Education or Outreach and a sizeable proportion of users are tertiary or school students and teachers. This demonstrates the excellent use of Ozboneviz to provide a novel means for the general public to access and interact with Australia’s museum collections.

The volume of downloads also emphasizes the usefulness of 3D digital service collections to the curating institutions that publish these data. Once digitised and published as ‘Open Download’, there is no further need for additional staffing or specimen handling, while the download volume far outstrips the capacity of collection managers to facilitate an equivalent rate of access to physical specimens. Because specimen downloads are monitored and usage is aggregated by MorphoSource, this expanded impact can be measured easily and is valuable for demonstrating the collection impact.

While the majority of the Ozboneviz collection came from our own digitisations, we incorporated data from other open-access MorphoSource collections – mostly from the oVert project collections network – to improve our coverage of species. This highlights the benefit of using the MorphoSource platform, which allows projects such as ours to derive new 3D imagery from published media in other collections and thus enhance the impact of an individual vouchered specimen (Blackburn et al. 2024). This can fill gaps in collections, as was the case for Ozboneviz, but it is also possible to create entirely new collections just from derivatives of other collection data. An example of this is the Fishboneviz collection, a sister project to Ozboneviz that relied purely on segmentation of already published CT scans to generate an 3D reference collection for Australian and Pacific Ocean fishes of zooarchaeological importance (Lambrides et al. 2024).

### Ozboneviz as a precedent for Australian open-access 3D biodiversity data

An important objective for Ozboneviz was the generation of a test case that would demonstrate the feasibility and clarify the challenges of open-access 3D biodiversity data in Australia. Our hope is that the success of this collection will inform a broader conversation about the value and management of open access FAIR 3D biodiversity imagery in the future. Many of the challenges we encountered reflect the issues raised in a submission by the National Imaging Facility Museums and Collections Special Interest Group to the ‘Accessing Australia’s Research Collections’ stakeholder consultation by the Australian Academy of Science (Weisbecker et al. 2024).

Appropriate archival storage for 3D imagery that also facilitates data discoverability and retrieval is currently an urgent challenge faced by curating institutions (Atlas of Living Australia et al. 2023). Large file sizes and secure archival storage have long been a problem for 3D imagery producers and users (Boyer et al. 2016, Rowe and Frank 2011). Collections managers are well versed in 3D imaging and recognise its value in collections preservation, data dissemination, research efficiency and public outreach (Hilton et al. 2021, Schindel and Cook 2018). Yet, these frontline managers cannot curate the large volumes of 3D imagery already being generated from their collections without institutionally supported storage and data management solutions (Weisbecker et al. 2024). This means these data, often acquired at high cost using competitive tax-payer funded grants, are effectively lost to the institution and wider research collection unless deposited on external data repositories (Davies et al. 2017, Hipsley and Sherratt 2019, Lewis 2019). The Ozboneviz initiative addressed this issue by identifying a digitisation strategy for a service collection that addressed a particular priority (in this case, zooarchaeology). This made it a sufficiently discrete work package to attract funding and addressed both the issue of storage and of high-quality presentation, with comparatively extensive longevity (the funding covers 14 years of MorphoSource storage and presentation). It is therefore a useful blueprint for further digitisation in Australia’s fragmented funding landscape.

It is important to note that the use of MorphoSource by Ozboneviz formalises an existing informal Australian trend of extensive deposition of 3D data. As of November 2024, excluding the Ozboneviz collection, 2598 3D media representing 2761 specimens held by Australian-based museums are currently stored on MorphoSource; 74% of these are published as open access; but in most cases, the managing organisation is not the collection from which the specimens were digitised. In establishing the Ozboneviz collection, we recognised that Australian collections managers are not yet resourced to manage these data but hope that it will demonstrate the feasibility of a 3D digital collections approach and thus encourage investment into the necessary infrastructure. Our goal is to transfer management of the Ozboneviz 3D imagery to the contributing institutions as soon as practicable.

### Opportunities, risks and limitations of implementing collections like Ozboneviz

Online repositories provide a technological solution to 3D data storage, but a range of concerns were frequently raised throughout our project. A major issue is the fact that both the storage and the presentation platform for Ozboneviz are overseas, in the US. There are valid concerns around the risks associated with the storage of digital specimen data on servers outside of Australia and external to the governance structures of the curating institutions. For example, many curating institutions that contributed specimens to Ozboneviz are state government agencies within Australia, but data uploaded to MorphoSource is subject to US laws (MorphoSource 2024). This could be overcome by generating Australian server space and even a separate Australian MorphoSource instance.

Another issue remains the assessment of risk in the provision of open-access 3D data, for example that 3D images might be monetised by private individuals (Matsui and Kimura 2022, Watanabe 2018). In the context of the large volume of 3D data generated by open-access initiatives such as oVert, this risk remains undemonstrated and may be negligible. However, as a means of managing such risks MorphoSource offers a range of options for data licensing, intellectual property assignment, and data management control.

Although these concerns are valid, it must be acknowledged that the process of making data more open and more FAIR will also likely coincide with some increased risk. We also highlight that 3D imagery are just one form of collections derived data. Within the context of Australian biodiversity collections there is ample precedent for accepting and managing such risks in the routine publication of other data modalities by collections such as occurrence data (Commonwealth Scientific and Industrial Research Organisation 2024) and two-dimensional photographic imagery (Australian National Botanic Gardens and Australian National Herbarium 2024) or genetic sequence data (Benson et al. 2015) by researchers. Ultimately, given the value of biodiversity collections in combating urgent global scale threats such as biodiversity loss, we argue that the risks are outweighed by the benefits of scientific capacity-building as our results already demonstrate.

Broadly, linkage and discoverability of collections derived imagery is another challenge in creating Australian 3D biodiversity service collections. For example, Ozboneviz imagery published to MorphoSource does not directly link to any platforms managed by the curating institutions or vice versa. Currently, MorphoSource can link digitised specimens to the US based iDigBio (2023) aggregator, but 3D specimen imagery is not yet discoverable via other key biodiversity data aggregators such as GBIF (2024) or the Atlas of Living Australia (Australian National Botanic Gardens and Australian National Herbarium 2024, Council of Heads of Australian Faunal Collections 2024). As with the provision of server space, this is an infrastructure issue that could be addressed by a separate, targeted initiative in the future to leverage the impact of 3D service and research collections. It is also worth highlighting that improving linkage and interoperability are core to initiatives such as the Digital Extended Specimen network (Hardisty et al. 2022) but the institutions that Ozboneviz collaborated with are not yet a part of such initiatives. However, MorphoSource is positioned well to grow these capabilities in the future through features such as the recently published MorphoSource Terms Vocabulary (MorphoSource 2024) which uses Darwin Core (Wieczorek et al. 2012) to reference specimens in collections in line with other repositories. As custodians of the source material for 3D biodiversity imagery, clear leadership, policies and guidelines from decision makers within curating institutions regarding imagery acquisition and data sharing will be crucial to establish an optimised and sustainable, open-access 3D data sharing culture in Australia.

It is also important to recognise that biodiversity collections represent the removal of plants and animals from Country and the custodianship of Indigenous peoples and is intimately tied to the extractive economics of colonisation and privileging of western epistemology and values (De Vos 2007, Mackenzie 2017, Weber 2021). The Ozboneviz collection does not and could not represent an attempt to rectify this situation. Nor do we suggest that access to 3D imagery should stand in lieu of repatriation of culturally significant objects to Indigenous communities affected by colonial collecting practices. Further investment and consultation are needed to ensure that FAIR 3D imagery also meets the CARE (Collective Benefit, Authority to Control, Responsibility, Ethics) principles for Indigenous data sovereignty (Carroll et al. 2020) to address the concerns and needs of Indigenous communities.

Our implementation of Ozboneviz has shown that service collections of 3D imagery are feasible and come with substantial benefits to the general and scientific public, giving them an important place in the future of Australia’s biodiversity collection sector. Our initiative has also identified clear limitations such as storage, presentation, and data security concerns, which all require separate, likely long-term, efforts to address. However, the successful roll-out of similar - and larger – initiatives in the US shows that these issues are surmountable. The use of MorphoSource storage and discrete service collections such as Ozboneviz are then a useful and safe avenue for managing the increasing data volumes in a best-practise way while suitable frameworks are being developed.

## Data Availability

All 3D imagery in the Ozboneviz collection is publicly available at https://www.morphosource.org/projects/000394988 under a CC-BY-NC 4.0 licence.

## Code Availability

No custom code was written for this paper.

## Acknowledgements

Ozboneviz was funded by the Australian Research Council Centre of Excellence for Australian Biodiversity and Heritage (CE170100015). Vera Weisbecker was additionally supported by an ARC Future Fellowship (FT180100634). We gratefully acknowledge the support of Duke University’s MorphoSource team – Doug Boyer, Mackenzie Shepard, Julie Winchester and Simon Choy. We thank the oVert team for support; Kirsti Abbott, Sue Horner and Sam Arman at the Museum and Art Gallery of the Northern Territory for facilitating collection access; Egon Perilli and Sophie Rapagna at the Flinders University Medical Device Research Institute for assistance and expertise in µCT imagery. Thanks also to Emma Sherratt and Toshiyuki Kimura for imagery and Nuttakorn Taewcharoen for segmentation assistance.

## Author Contributions

VW designed and supervised the project. EM acquired and uploaded 3D imagery and managed the project database. VW & EM co-wrote the manuscript and curated contributions. JvZ and PV managed the 3D imagery acquisition and upload process and acquired and uploaded 3D imagery. DF and JR acquired and uploaded 3D imagery. SI, TM, KM, SOC, TP, SSC, DS, AY provided advice on specimen and skeletal element selection and facilitated access to biological collections. AL contributed to databasing protocols and supported the stakeholder consultations. All authors have edited sections of the paper according to their expertise.

## Supplementary Methods

The Ozboneviz digitisation initiative was a collaboration of a network of stakeholders. It consists of a core team of project managers and digitisation officers, as well as contributing staff within collections who facilitated access, co-acquired data, and supported the process of acquiring permissions. The core team included the project lead (Vera Weisbecker), several academic stakeholder-advisors, the equivalent of one year’s full-time position of a project manager and chief acquisition officer (Pietro Viacava followed by Jacob van Zoelen), supported by several casual and short-term project officers tasked with database management, 3D image acquisition, segmentation and data upload (Erin Mein, Diana Fusco and Jorgo Ristevski). Funding for these positions was provided by the Australian Research Council Centre of Excellence for Australian Biodiversity and Heritage (CABAH) through a Legacy Grant Scheme.

In consultation with stakeholders and project collaborators, we developed a target list of 130 Australian mammals, 28 New Guinean mammals, 11 birds, 36 reptiles and 5 frogs. We prioritised digitising the skull and eight major elements from the appendicular skeleton for each species. Mammalian and reptilian postcranial bones consisted of the scapula, humerus, ulna, pelvis, femur, tibia, astragalus (talus) and calcaneum (calcaneus) (Figure 1 of the manuscript). For birds we digitised eight bones including the coracoid-scapula, carpometacarpus, tibiotarsus and tarsometatarsus. For frogs we aimed to digitise 10 bones including the coracoid and calcaneum. Skeletal elements such as vertebrae, phalanges and ribs which are numerous and therefore, less optimal for estimating minimum numbers of individuals in palaeozoological contexts, were not digitised. In the majority of cases, we chose not to digitise the radius or fibula, although some exceptions included bats (Chiroptera), birds (Aves), frogs (Anura) and artiodactyls (Artiodactyla) where these elements are either taxonomically distinctive or fused to the ulna or tibia. Specimens were digitised from a range of museums (a breakdown is in Table 1).

**Table 1.** Number of specimens digitised by curating institution. *CT imaged specimens sourced from oVert collaborating institutions via MorphoSource.

| Curating Institution | No. Specimens |
| --- | --- |
| South Australian Museum | 121 |
| Australian Museum | 68 |
| Museum and Art Gallery of the Northern Territory | 38 |
| The University of Queensland | 19 |
| The Australian National University | 7 |
| Australian National Wildlife Collection | 6 |
| Flinders University Palaeontology Laboratory | 6 |
| California Academy of Sciences* | 2 |
| Laboratory of Emma Sherratt, University of Adelaide | 2 |
| Michigan State University Museum | 2 |
| University of Florida, Florida Museum of Natural History* | 2 |
| Harvard University Museum of Comparative Zoology* | 1 |
| Museum of Vertebrate Zoology, UC Berkeley* | 1 |
| Museum of Zoology, University of Cambridge | 1 |

### Biodiversity collections

We digitised vouchered specimens from Australian-based biodiversity collections at the South Australian Museum, Museum and Art Gallery of the Northern Territory, the Australian Museum and the Australian National Wildlife Collection. Specimens from these institutions were prioritised as each have clearly defined public access protocols and specimen metadata is rich, formatted to Darwin Core standards and discoverable via biodiversity data aggregators such as OZCAM (Council of Heads of Australian Faunal Collections, 2024), GBIF (GBIF 2024) and iDigBio (iDigBio 2023).

Secondary sources of specimens were the University of Queensland Archaeology Laboratories, the Australian National University Zooarchaeology Collection and Flinders University Palaeontology Laboratory. Select specimens that were difficult to source were segmented from computed tomography image stacks deposited on MorphoSource.org by international institutions or private collections. We also received donations of 3D imagery of cetaceans from Dr Toshiyuki Kimura of the Gunma Museum of Natural History (Kimura 2023).

### Imaging modalities

We used three 3D imaging modalities, structured light surface scanning, micro-computed tomography (µCT) and photogrammetry. All µCT was undertaken at the Flinders University Medical Device Research Institute using a Nikon XT H 225ST. As our budget for µCT and subsequent mesh segmentation was limited, we prioritised surface scanning dry specimens. Structured light scanners were brought to each collection to digitise specimens in-house rather than initiate large numbers of specimen loans. We used multiple structured light surface scanning devices consisting of a Polyga Compact S1, a Solutionix C500, a Shining 3D Einscan Pro+ and an Artec Space Spider. As each has optimal operating conditions the choice of device was determined on a balance of specimen size, collection location, light conditions and project stage. For example, we used the Polyga (point distance = 0.085mm) for very small specimens and the Einscan Pro+ (point distance = 0.24mm with turntable and 0.7mm when handheld) for medium to large specimens in low-light conditions. We acquired the Artec Spider (point distance = 0.15mm) part-way through our digitisation program after which it became the preferred device for digitising medium and large specimens. The Solutionix was located at the University of Queensland and used for digitising specimens from the Archaeology Laboratories. Texture (photorealistic colour) capture was not a priority and not available across all devices but, where possible, was captured to increase the utility of the 3D imagery. Photogrammetry using Agisoft software was only used for few large specimens when the available surface scanning devices were unsuitable or unavailable at the time of scanning.

### Specimen selection

Specimens were selected for digitisation based on a balance between metadata completeness, specimen condition and species rarity. Because our goal was to create a service collection for researchers, it was important that end users could account for potential influences of animal age, sexual dimorphism or geography on skeletal morphology. We therefore aimed to digitise adult, wild specimens that were free from pathologies with metadata on sex and collection location. However, we found the completeness of these metadata varied widely owing to historical realities of collections management. When choosing suitable specimens, we avoided clearly juvenile specimens based on molar eruption and fusion of epiphyses. Known captive specimens (e.g. from zoos) were also avoided when possible, owing to their potential for morphological differences with wild specimens (e.g. Crossley and del Mar Miguélez 2001, Hartstone-Rose et al. 2014, Mitchell et al. 2021). We digitised captive animals only as a last resort after we had exhausted other avenues to source wild specimens. Type specimens were not specifically targeted for digitisation as many are held in collections outside of Australia but, were treated as high priority where available.

Computed tomography was used for particularly rare and important species. This presented its own challenges; in some cases, metadata did not record details on specimen sampling and preparation methods that can impact on digital imaging. For example, roadkill specimens can have substantial damage to the skeleton that is unable to be ascertained prior to µCT (Figure 1). Historical specimens of bats and birds were once commonly collected via shooting. In the New Guinea naked-backed fruit bat specimen we found embedded lead shot pellets that produced substantial noise in the computed tomography images and made the segmentation of individual bones difficult. In some cases, identifying where the skull of fluid-preserved specimens had been removed and replaced with foam formers was difficult and resulted in additional imaging costs and time spent by our team and curatorial staff to identify and reimage the head separately (such as the extremely rare marsupial mole *Notoryctes caurinus*) or more appropriate specimens. However, the reuse value of the µCT volume stacks of these ‘headless’ or damaged specimens is high, and we chose to make these data available on MorphoSource as well. These instances highlight the vital importance of metadata discoverability to collections users and the need to resource curatorial staff with the appropriate tools that allow for specimen lifecycle traceability and digitally catalogued links between all data points of the Digital Extended Specimen (Hardisty et al. 2022, Lendemer et al. 2020, Webster 2017).

**Figure 1.**
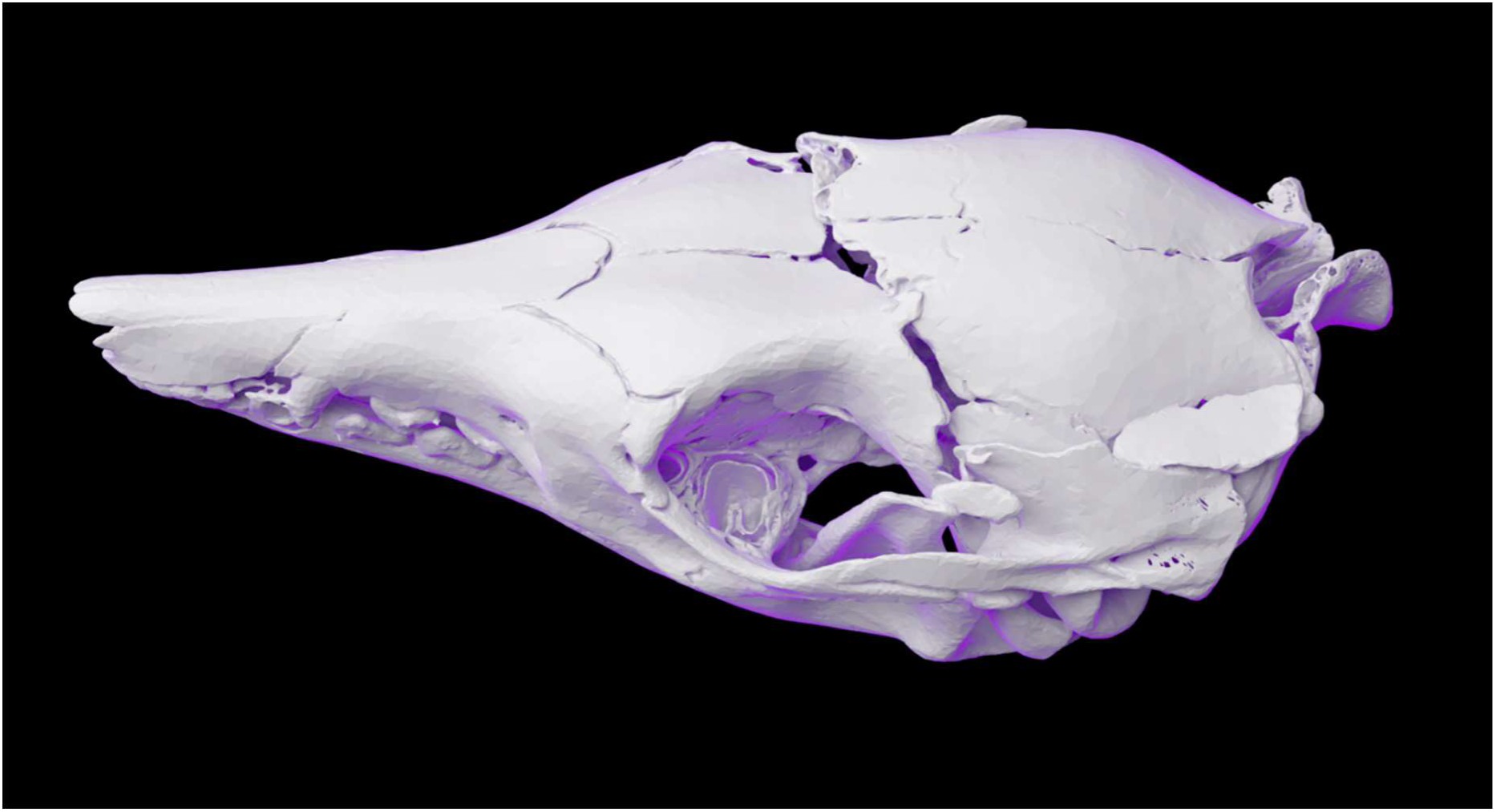
Damaged skull of a greater bilby (*Macrotis lagotis*) revealed after µCT imaging. This specimen (SAMA-M3600) was unsuitable for creating 3D models for the Ozboneviz collection but the µCT volume stacks are made available on MorphoSource.org.

### 3D image generation

3D point clouds acquired via structured light scanners were meshed using the proprietary software associated with each scanning device. This included Flexscan3D (Polyga 2021) ezScan2017 (Solutionix) (Medit 2019), Einscan Pro v2.6.0 (Shining 2022) and Artec Studio 17 Pro (Artec 2022). The 10 target skeletal elements were segmented from µCT volumetric image stacks using Materialise Mimics v. 24 (Materialise 2022) and 3D Slicer (Fedorov et al. 2012). We used Agisoft Metashape v.1.8.2 (Agisoft 2022) to generate 3D meshes from digital photographs. 3D meshes were exported as .PLY format files or Wavefront .OBJ files if texture was captured. We avoided making extensive modifications to the exported meshes but in some cases filled small holes or smoothed meshes in Geomagic Wrap v2021.2.2 (3D Systems 2021) while noting these modifications in the file metadata.

### Data sharing

We purchased storage on the MorphoSource.org platform because it has been designed as a FAIR 3D data archive that balances the needs of data producers, users and curating institutions (Boyer et al. 2016). The architecture of the MorphoSource repository allows for connections between 3D data and specimen metadata to be maintained, as well as supporting metadata describing imagery acquisition, processing, intellectual property and copyright. MorphoSource also has the capacity for data managers and curating institutions to authorise user access to 3D data and track data downloads, intended usage and user demographics. We uploaded 3D meshes derived from our three imaging modalities to MorphoSource alongside digital photographs of the physical specimen and any catalogue tags or accompanying documentation to maximise information on specimen condition and provenance. We also uploaded all volumetric image stacks produced via µCT and linked these as ‘Parent Media’ to the 3D meshes of that specimen derived by segmentation. We chose not to upload raw point cloud data owing to the large file sizes and dependence on proprietary software which makes these poor candidates for data archiving. At a minimum all 3D imagery is provenanced to a physical specimen using Darwin Core (Wieczorek et al. 2012) standard terminology for institution, collection, catalogue number and taxonomy and is further described by skeletal element and body side. Where it exists, each 3D image was also linked to a corresponding occurrence record on the iDigBio (iDigBio 2023) biodiversity data aggregator. Although iDigBio ingests data from the Atlas of Living Australia, the facility to link 3D imagery on MorphoSource directly to this Australian based data aggregator is not yet available.

Where taxonomic names imported from iDigBio clash with more recent specimen identifications or taxonomic revisions we made use of the MorphoSource facility to include both taxonomies. All 3D imagery in the Ozboneviz collection has metadata describing imaging modality, device, operator, post-processing interventions that have altered the mesh surface and descriptions of any conditions that may obscure or alter the natural bone morphology such as adhering soft tissue, damage or scanning artefacts. All 3D imagery deposited on MorphoSource are assigned an Archival Resource Key (ARK) as a persistent identifier. MorphoSource also have the capacity to mint Digital Object Identifiers (DOI) and we strongly encourage researchers to request and cite DOI’s for each 3D media they use.

Intellectual property over the 3D imagery is assigned to the curating institutions (i.e. museum or university). All 3D imagery produced by Ozboneviz is provided to users as ‘Open Download’ under a Creative Commons BY-NC 4.0 copyright licence which permits non-commercial use, sharing and adaptation of these data.

